# When the brightest is not the best: illuminant estimation from the geometry of specular highlights

**DOI:** 10.64898/2026.01.22.700600

**Authors:** Takuma Morimoto, Robert J. Lee, Hannah E. Smithson

**Affiliations:** Department of Experimental Psychology, University of Oxford, Oxford, UK; Cambridge Research Systems Ltd

**Keywords:** Colour constancy, illuminant estimation, specular highlights, brightest-is-white heuristic

## Abstract

Colour constancy supports stable object-colour perception across changes in illumination. Illuminant colour can be inferred from white surfaces or specular highlights, and many models adopt a “brightest is white” heuristic to identify illuminant colour. We tested an alternative: observers use structured changes in the proximal image to locate highlights even when they are not the brightest elements. Using computer-rendered scenes, we manipulated the reliability of two cues—brightest element and highlight geometry. Each scene showed a single textured sphere lit by multiple point lights with identical spectra. The sphere had uniform spectral reflectance, while a noise texture attenuated the reflectance by a variable scale factor. We tested three specularity levels (matte, low, mid). Observers viewed a 1.5 s animation and judged whether chromatic changes reflected an illuminant or material change. Performance for matte surfaces was near chance but improved sharply with increasing specularity. Observers outperformed an ideal observer relying solely on the brightest element. When highlights landed on darker texture regions, performance improved further, contrary to the brightest-element prediction. Phase scrambling, which disrupted highlight geometry while preserving global statistics, substantially reduced performance. Overall, observers do not rely solely on the brightest element; they exploit diffuse–specular regularities to resolve surface–illuminant ambiguity.

## 1. Introduction

The way light interacts with an object provides the visual system with insights into the object’s properties. For example, the spectral reflectance of an object, determined by its pigments, can provide biologically important details such as fruit ripeness [1]. More complex optical interactions, such as transmission and specular (mirror-like) reflection, produce sensations of transparency [2] and glossiness [3, 4]. Spatial variations in colour and luminance across an object’s surface can also reveal material classes (e.g., plastic, ceramic, rubber) [5] and physical states like wetness [6]. Furthermore, many organisms—including plants and insects—have evolved to exploit specific colour patterns and light interactions for camouflage or mimicry, helping them avoid detection by predators or to make themselves conspicuous for reproductive advantage [7–10]. However, reliable extraction of surface properties is challenging primarily because the proximal image of an object is contaminated by surrounding lighting conditions and is dependent on viewing geometry. There are indications from the natural world that mimicry can include not only the colour and patterning of an object, but also the geometry of diffuse and specular components of light-material interactions: a moth, Eudocima aurantia, successfully renders a 3D surface on a flat wing using nanostructures to control both the directionality and spectrum of the reflected light [11]. This example, in which the characteristics of diffuse and specular reflections are embodied in animal morphology, suggests that the complexities of light-material interactions are widely sensed.

This study investigates how humans separate lighting and material components in the context of colour constancy—the phenomenon where object colours appear consistent despite changes in the spectral content of the lighting. Color constancy has been extensively studied, with research spanning behavioral experiments [12,13] and computer vision algorithms designed to adjust white balance in images [14–17]. However, many previous experimental studies have largely relied on simplified stimuli, typically featuring flat, matte surfaces. Although the field has increasingly moved towards more complex and naturalistic stimuli and environments [18–22], the contributions of different stimulus features to colour constancy remain relatively unknown. Here, we demonstrate how introducing material complexity into experimental scenes can challenge widely-known visual strategies for inferring illumination colour.

In theory, the colour of the illumination could be directly accessed if a scene contained a white surface with a spectrally flat reflectance because such a surface would reflect the illumination without altering its spectral properties. Similarly, real-world materials often exhibit specular reflections that return the light spectrum unchanged [23–33]. A useful property of white surfaces and of specular highlights is that they are often the brightest element in a scene. Thus, by assuming that the brightest element is either a white surface or a specular highlight, we can identify a chromatic signal that provides the closest representation of the scene’s illumination. This heuristic is known as the “brightest is white” assumption [34], and it has explained empirical data in early colour constancy studies that employed matte surfaces [35–40].

However, this heuristic is not always valid; for example, when a glossy object has a high-contrast texture, the specular highlight may fall on a dark region of the surface, meaning that the centre of the highlight is not necessarily the brightest point. This is illustrated in Figure 1. In panel *a*, the specular highlight appears on a bright part of the texture, making the brightest point and the highlight centre spatially coincide (spatially-aligned condition). In panel *b*, the specular highlight falls on the dark part of the texture, and the brightest point does not align with the centre of the specular highlight (spatially-separated condition). Chromaticity diagrams [41] on the right illustrate why the “brightest is white” assumption is not the most informative. The plus symbol represents the chromaticity of the diffuse component, while the black cross indicates the chromaticity of the specular component alone—corresponding to the illuminant’s chromaticity, which we aim to infer. According to the Ward reflectance model [42], which provides an adequate physical description of the light-material interactions relevant to our stimuli, the light reflected from any point on the surface is simply a weighted sum of diffuse and specular components where the weights vary over the object’s surface. Thus, the object’s “locus of chromaticities” follows a straight line between the plus and cross symbols. The magenta, cyan, and green symbols denote the locations of three representative points on the surface and their corresponding chromaticities. In both conditions, the green square has the chromaticity furthest from the illuminant because it comes from a region with only diffuse reflection and no specular component. In the spatially-aligned condition, both the magenta diamond (brightest point) and the cyan circle (specular highlight centre) provide the best information about the illuminant. In the spatially separated condition, the cyan circle is more informative than the magenta diamond, because it is less contaminated by the diffuse component. In this case, the “brightest element” heuristic picks the magenta diamond because its overall intensity is higher than that of the highlight’s centre. Based on this observation, we predict that observers will demonstrate greater colour constancy than the brightest element assumption suggests—if they can use regularities in features of the specular highlight, such as its geometry, and consistently extract the illuminant estimate from regions of the image that are dominated by specular reflection. This study directly tested whether the violation of the “brightest element” heuristics impairs performance in an operational colour constancy task. Here, ‘operational colour constancy’ refers to observers’ ability to discriminate an illuminant change from a reflectance (material) change, rather than to judge or match surface colour directly [43]. It is a performance-based measure, rather than an appearance measure.

**Figure 1:**
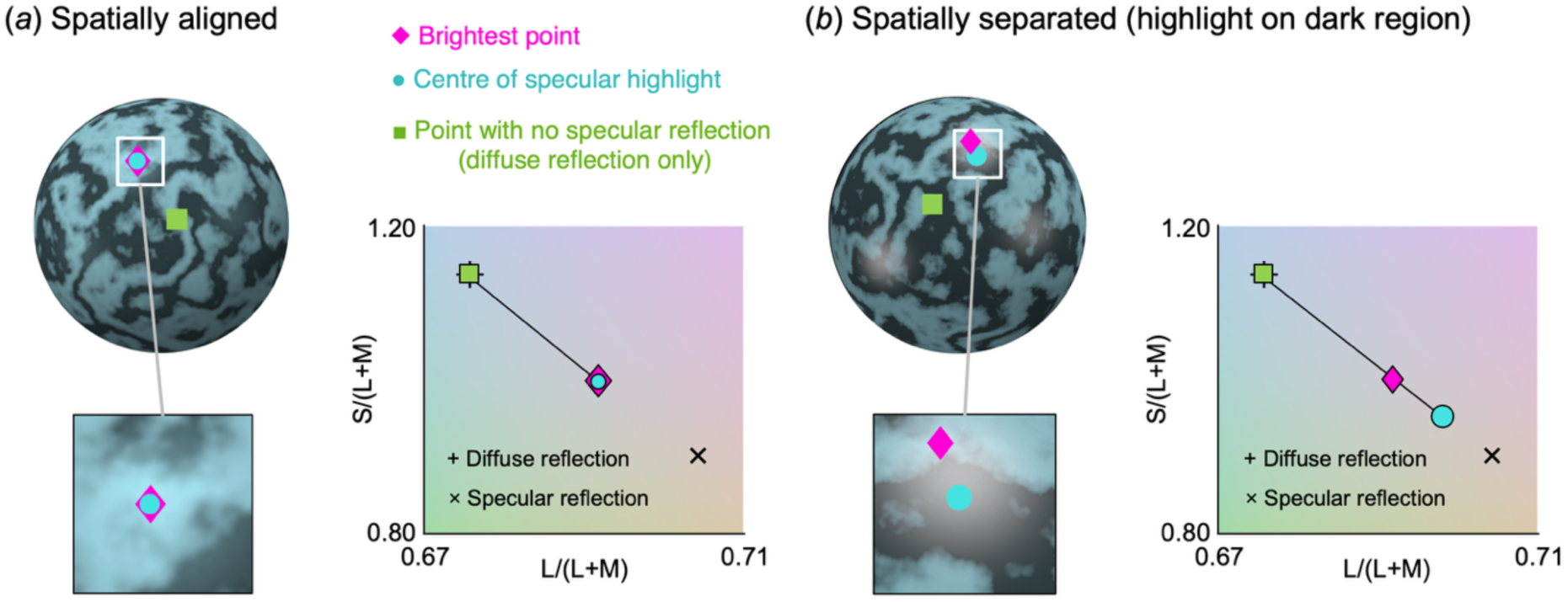
Manipulation of the reliability of the “brightest element” heuristic by positioning specular highlights on either bright or dark regions of a high-contrast texture. See main text for details.

## 2. Method

### (a) Apparatus

The experiment was conducted in a dark room. Stimuli were presented on a CRT monitor (53.3 centimetres, 1024×768 pixels, NEC, FP2141SB, Tokyo, Japan) controlled by a visual stimulus generator (ViSaGe MkII, Cambridge Research Systems, Rochester, UK), which allows 14-bit intensity resolution over full dynamic range for each phosphor. Gamma correction was performed using data from a colorimeter (ColorCAL MK II, Cambridge Research Systems, Rochester, UK) and spectral calibration was conducted with a spectroradiometer (SpectroCAL MKII, Cambridge Research Systems, Rochester, UK). Viewing distance was maintained by a chin rest positioned 92 cm from the CRT monitor.

### (b) Rendering

All stimuli were generated by computer graphics techniques. The geometry of each scene, i.e. location of the viewpoint or camera, an object and three point-light sources, was defined using the 3-D modelling software, Blender 2.78b (Blender Foundation, Amsterdam) and, rendering was conducted using the physically-based renderer, PBRT (pbrt-v2, Physically Based Rendering Tool, [44]). The resultant multispectral images (31 channels, from 400 nm to 700 nm with 10 nm steps) were converted to LMS cone coordinates based on the Stockman and Sharpe 2° cone fundamentals [45, 46] and then finally converted to RGB values for the stimulus presentation. We used MATLAB 2016b (MATLAB 2016b, Mathworks, Massachusetts, USA) and RenderToolbox3 [47] to automate the stimulus generation process.

### (c) Stimuli

We used smooth spheres of radius 10.1 degrees of visual angle that had a constant roughness of 0.02 defined in PBRT. In PBRT, roughness parameterises the microfacet distribution of the surface: lower values concentrate the specular lobe into a tighter, sharper highlight, whereas higher values spread the reflected light over a wider range of directions, giving a broader, more diffuse highlight. We used a low value (0.02) that was fixed for all stimuli so that highlights remained compact and well localised. Each scene contained only a single sphere illuminated by three point sources of light, each with the same spectral content. The positions of the three point sources were set manually. They were placed behind the viewpoint such that the resulting highlights were clearly visible, spatially well separated, and distributed across a range of texture positions. We used ten distinct configurations to reduce the possibility of observers memorising fixed highlight locations. These ten configurations specified the light-source positions only; whether a given trial was spatially-aligned or spatially-separated was determined by the texture manipulation (see below), not by the lighting configuration itself. The sphere had a single spectral surface reflectance, but across the surface of the sphere, the spectral reflectance was attenuated by a wavelength-independent scalar variable in a location-specific manner, giving rise to a spatial texture. The textures were achromatic high-contrast Perlin noise patterns (random scalar reflectance maps with values between 0.2 and 1.0), with ten patterns generated from different random seeds. The spatially-aligned and spatially-separated conditions were controlled by rotating the texture in the image plane about the image centre. In the spatially-aligned condition the most intense highlight region coincided with a bright texture region, so that the brightest image point and the highlight centre coincided; in the spatially-separated condition the highlight fell on a dark region, so that the brightest point and the highlight centre did not coincide (Figure 1). We used three specular levels: 0 (matte), 0.02 (low), and 0.04 (mid), corresponding to 0%, 2%, and 4% specular reflectance. Specular reflectance was modelled using the Ward reflectance model [42]; details are given in the Supplementary Information. To investigate the role of specular highlight geometry, we also included phase-scrambled images that preserved the original image’s chromatic and luminance distribution but disrupted its spatial structure.

To quantify the observer’s performance at discriminating a reflectance change from an illuminant change, we needed to prepare a sufficient set of animations for both scenarios. First, we chose two illuminations, sunlight and skylight (Figure 2*a*). Second, we chose 240 publicly available reflectance spectra of natural and man-made objects [48–50]. For illuminant-change animations, the chromaticity shift caused by changing from sunlight to skylight, or vice versa, falls along a yellow–blue or blue–yellow axis (panel b, left). For reflectance-change animations, by contrast, there is no corresponding constraint. With arbitrary choice of reflectance pairs, the chromaticity change could be in any colour direction. To prevent the axis of chromaticity change being a reliable cue separating reflectance-change versus illuminant-change, we selected the reflectance pairs so that the direction of their chromaticity change matched the sunlight–skylight axis. Specifically, we computed the chromaticity shift produced by candidate reflectance pairs and retained 50 pairs whose shift direction lay close to this axis (panel b, right).

**Figure 2:**
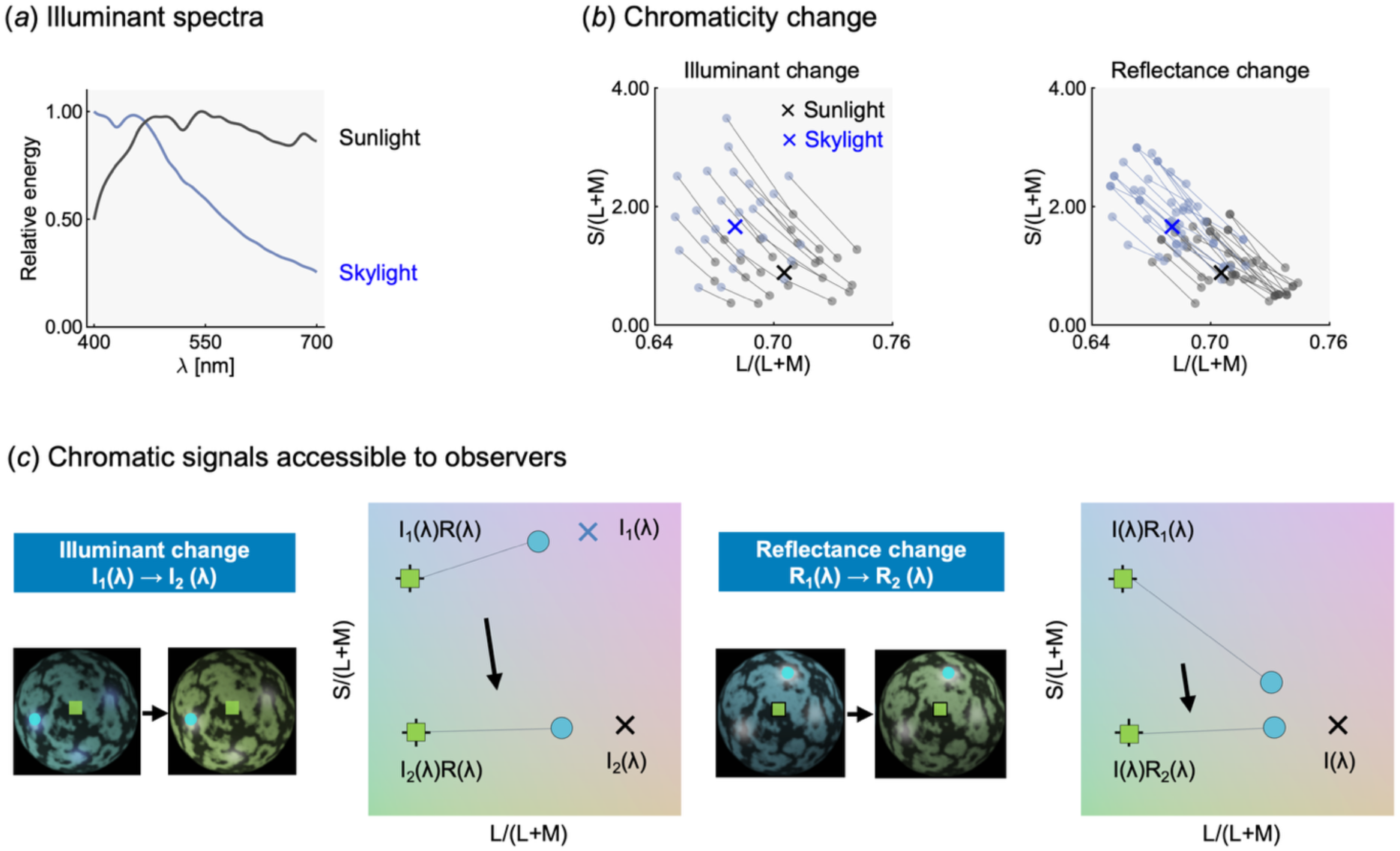
Spectra of the illuminants and chromatic signals for illuminant vs. reflectance changes. (*a*) Spectral compositions of sunlight and skylight. (b) Plots show the chromatic change of the diffuse component of the reflected light (not the diffuse reflectance of the object) for the illuminant change condition (left) and the reflectance change condition (right). For the illuminant change, each pair of symbols (blue under skylight and grey under sunlight) represents one reflectance. For reflectance change, the grey lines represent changes under sunlight, while the blue lines represent changes under skylight. The chromatic change directions are similar in both cases. (*c*) Chromatic signals available in illuminant-change and reflectance-change trials. Since reflectance pairs were chosen such that the distribution of chromatic change in the diffuse component was approximately matched for reflectance change and illuminant change trials (green squares), specular signals are critical for the task (cyan circles).

Panel *c* depicts example trials for both conditions, describing the change of chromatic distribution during animation: an illuminant change from skylight *I*_1_(*λ*) to sunlight *I*_2_(*λ*) (left plot) and reflectance change from *R*_1_(*λ*) and *R*_2_(*λ*) (right plot).

### (d) Observers and Ethics

Six observers (four female, two male: AKH, TD, JH, SR, TM, and HH) participated in all conditions. The observers had an age of 25.8 ± 4.17 (mean ± 1 SD) and all had normal colour vision, as confirmed by Ishihara or Hardy-Rand-Rittler pseudoisochromatic colour plates. AKH and SR had previously participated in related experiments with the same task [31] but were unaware of this study’s purpose. JH, TD, and HH were naive observers. TM is the first author. The study was approved in 2016 by the Medical Sciences Inter-divisional Research Ethics Committee at the University of Oxford in agreement with the Declaration of Helsinki (6th revision, 2008). All observers gave written informed consent before participating.

### (e) Procedure

In each trial, observers watched a 1.5-second animation consisting of 10 frames, during which the image of a sphere gradually changed colour. Their task was to determine whether the colour change resulted from a change in illuminant colour or a change in surface colour [30–32, 51–54]. Immediate feedback was provided after the response. Each session comprised 500 trials: 100 trials for each specular level (zero, low, and mid) in the original sphere condition, and 100 trials for two specular levels (low and mid) in the phase-scrambled condition. Within each specular level, half of the trials corresponded to reflectance changes, while the other half corresponded to illuminant changes. Furthermore, half of the trials were the spatially-aligned condition (i.e. the most intense highlight region fell on a bright texture region), while the remaining trials were the spatially-separated condition (i.e. the highlight fell on a dark texture region.) Before the main experiment, all observers completed a block of 100 practice trials (original, non-scrambled sphere stimuli, presented with trial-by-trial feedback). Six observers completed four sessions, totaling 2 000 trials, with a break after every 125 trials. The order of conditions was randomised for each observer.

### (f) Simulation using an ideal observer model

We simulated an ideal observer model that bases its judgments about material or illumination changes on the chromatic changes in regions or statistical features of interest (e.g., the brightest pixel). The model follows a three-step process: observing an animation and extracting relevant signals (Figure 3*a*), computing chromatic distance (Figure 3*b*), and learning a probability density distribution as a function of chromatic change for each classification category (Figure 3*c*). Once the probability function is developed, the model classifies new trials based on observed values.

**Figure 3:**
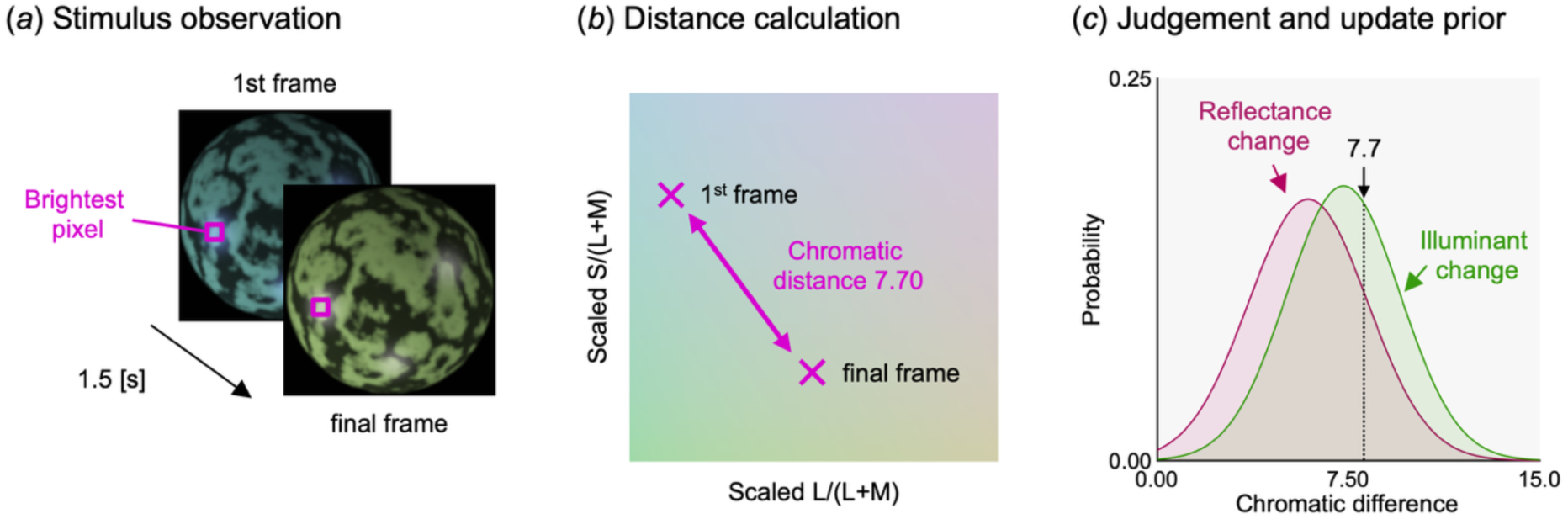
Schematic of a simulation for a specific visual strategy. (*a*) The model extracts chromaticity from a point of interest (e.g. brightest pixel) in both the first and final frames. (*b*) The model then computes the chromatic distance between frames using the MacLeod-Boynton chromaticity diagram, where axes are scaled by detection thresholds to approximately equate perceptual distances. (*c*) Based on the observed chromatic difference, the model classifies the trial and updates the prior distribution after each trial integrating new observations.

We implemented a naive Bayes classifier using the fitcnb function from Statistics and Machine Learning Toolbox in MATLAB. We analysed only the first and final frames, as their chromatic change provided the most informative signal. In brief, a naive Bayes classifier estimates, separately for each category (illuminant change versus reflectance change), the probability distribution of the observed chromatic change; a new trial is then assigned to whichever category yields the higher posterior probability for its observed chromatic change. Chromatic distance was computed in a scaled MacLeod-Boynton (MB) chromaticity diagram, where equal-energy white is the origin. Each axis was independently scaled by chromatic discrimination thresholds along L/(L+M) and S/(L+M) axes, measured before the experiment using 2-degree homogeneous circular colour patches. These thresholds were obtained from two observers in a separate preliminary session using a four-alternative forced-choice (4AFC) odd-one-out staircase procedure. On each trial, four patches, each 2 deg in diameter, were presented in a 2 × 2 array: three showed the mid-grey reference and one was a chromatically displaced target. Observers reported the odd one out, and the chromatic distance of the target from mid-grey was varied by a QUEST adaptive staircase [55] to estimate the discrimination threshold along the L/(L+M) and S/(L+M) axes.

During the training phase, the classifier observed the same set of 100 practice trials under the original sphere condition that human observers completed before the main experiment. It separately compiled statistics for reflectance and illuminant change trials based on the distributions of observed chromatic changes in the statistics of interest. No prior information was given before training.

In the test phase, the classifier processed all trials from the main experiment. For each trial, it computed the chromatic difference and classified it as more likely belonging to the reflectance or illuminant change distribution. After each trial, the prior distribution was updated for the next classification.

## 3. Results

### (a) Sensitivity estimate *d’* and response bias estimate *C*

We used a bias-free estimator of sensitivity (*d’*) to evaluate the classification performance between illuminant and reflectance changes, and criterion (*C*) as an estimate of response bias where positive values indicate a bias toward classifying a trial as an illuminant change. Figure 4*a* shows *d’* for individual observers. For all observers, as predicted, at zero specularity, performance was near chance level (*d’* = 0), but it improved as specularity increased. Notably, when specular highlights appeared on darker regions of the texture (the spatially-separated condition), observers’ performance further improved. Another key finding is that performance dropped significantly when the spatial structure of the specular highlight was disrupted (panel *b*), suggesting that the highlight’s geometry plays a critical role in distinguishing reflectance from illuminant changes. Due to high consistency across observers, the average value across observers provides a representative summary (panel *c*). Panel *d* shows *C* values, which remained close to zero, indicating no strong response bias toward either category.

**Figure 4:**
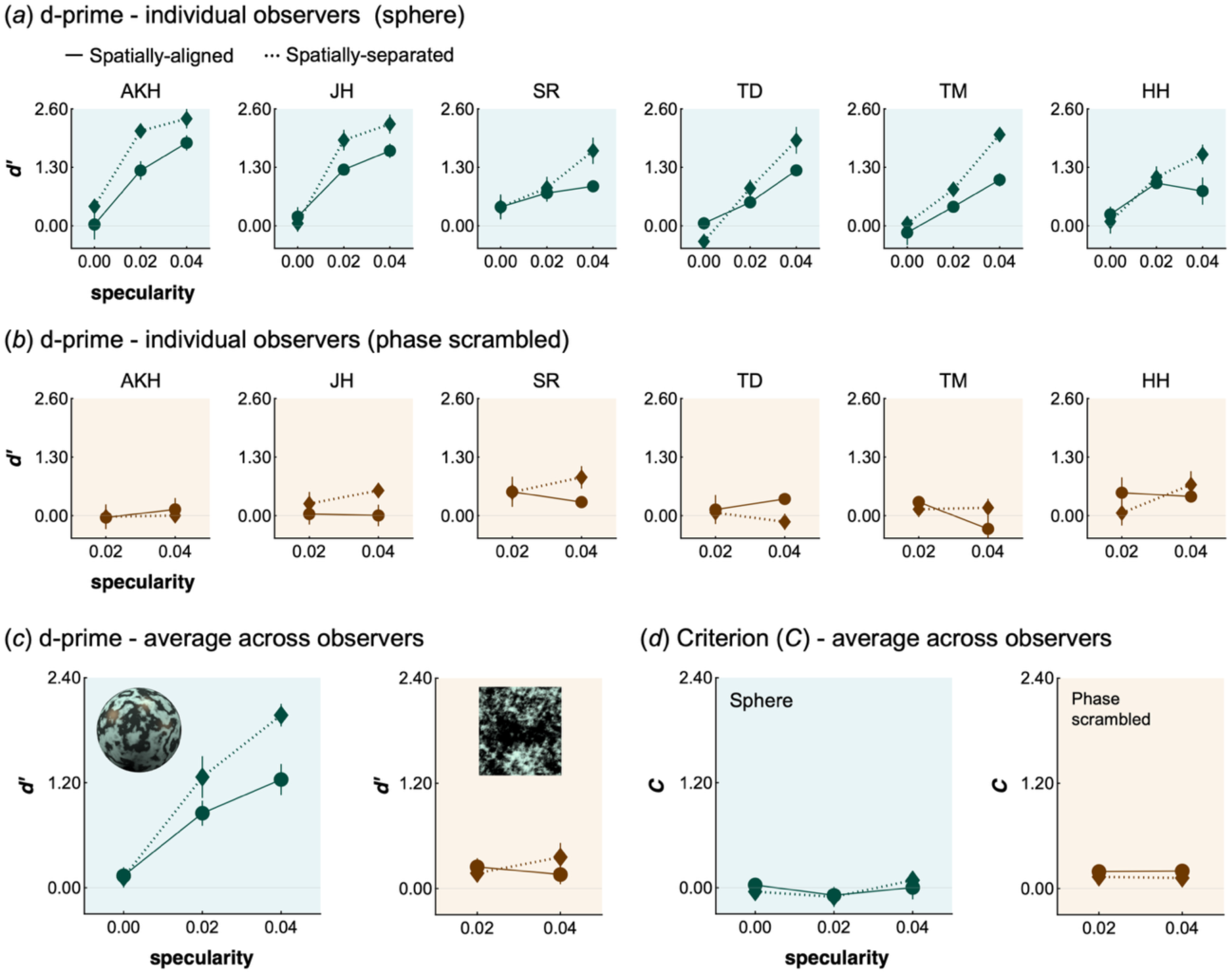
(*a*) Discrimination sensitivity (*d’*) estimates for the sphere condition, shown for six individual observers. The solid line represents *d’* for spatially aligned trials, while the dashed line represents spatially separated conditions. Values are averaged across four sessions, with error bars indicating the standard error across sessions. (*b*) *d’* in the phase-scrambled condition. Values and error bars are consistent with those in panel a. (*c*) Averaged *d’* across six observers for sphere condition (left) and phase scrambled condition (right). Error bars show standard error across observers (n=6). (*d*) Criterion *C* where positive values indicate a greater tendency to respond to illuminant change. Error bars show standard error across six observers.

To statistically confirm the effect, we performed two-way repeated-measures analysis of variance (ANOVA) on the average *d’* in the original sphere condition (left plot in panel c), with spatial alignment (spatially aligned and spatially separated) and specularity (zero, low and mid) as within-subject factors. Significant main effects were found for spatial alignment (*F*(1,5) = 33.0, *η*^2^ = 0.868, *p* = 0.00225) and specularity (*F*(2,10) = 43.2, *η*^2^ = 0.896, *p* < 0.001) as well as a significant interaction (*F*(2,10) = 13.81, *η*^2^ = 0.734, *p* = 0.00133). Further analysis revealed no effect of spatial alignment at specularity zero (*F*(1,5) = 0.03, *p* = 0.869) but significant effects at low and mid (*F*(1,5) = 11.3, *p* = 0.020074; *F*(1,5) = 111.2, *p* = .000132). Specularity was significant for both spatially-separated and aligned trials (*F*(2,10) = 21.8, *p* < 0.000226; *F*(2,10) = 52.4, *p* < 0.001). Bonferroni corrections showed significant differences in the spatially-separated condition between zero and low, zero and mid, and low and mid. For the spatially-aligned condition, differences were significant between zero and low, and zero and mid.

For the phase scrambled image (right panel in panel c), a two-way repeated-measures ANOVA with specularity levels low and mid revealed no significant main effect of spatial alignment (*F*(1,5) = 0.36, *η*^2^ = 0.0680, *p* = 0.575), specularity (*F*(1,5) = 0.41, *η*^2^ = 0.0757, *p* = 0.550), or their interaction (*F*(1,5) = 2.01, *η*^2^ = 0.287, *p* = 0.215).

### (b) Simulating “brightest element” and “centre of specular highlight” heuristics

Figure 5 illustrates the performance of computational observers using the “brightest element” heuristic (panel *a*) and the “centre of specular highlight” strategy (panel *b*). For the latter, each animation contained three specular highlights, leading to the construction of three separate models, each focusing on the most intense, second most intense, or third most intense highlight (1st, 2nd, and 3rd, respectively). The “brightest element” heuristic produced results that differed qualitatively from those of human observers: whereas human d′ increased in the spatially-separated condition, the brightest-element model showed the opposite trend, its performance falling when the highlight moved onto a dark region. In this configuration, the brightest point identified a location that carried stronger diffuse contamination. The “centre of specular highlight” model instead showed the human-like pattern, improving in the spatially-separated condition, since now the model located a specular reflection with little diffuse contamination.

**Figure 5:**
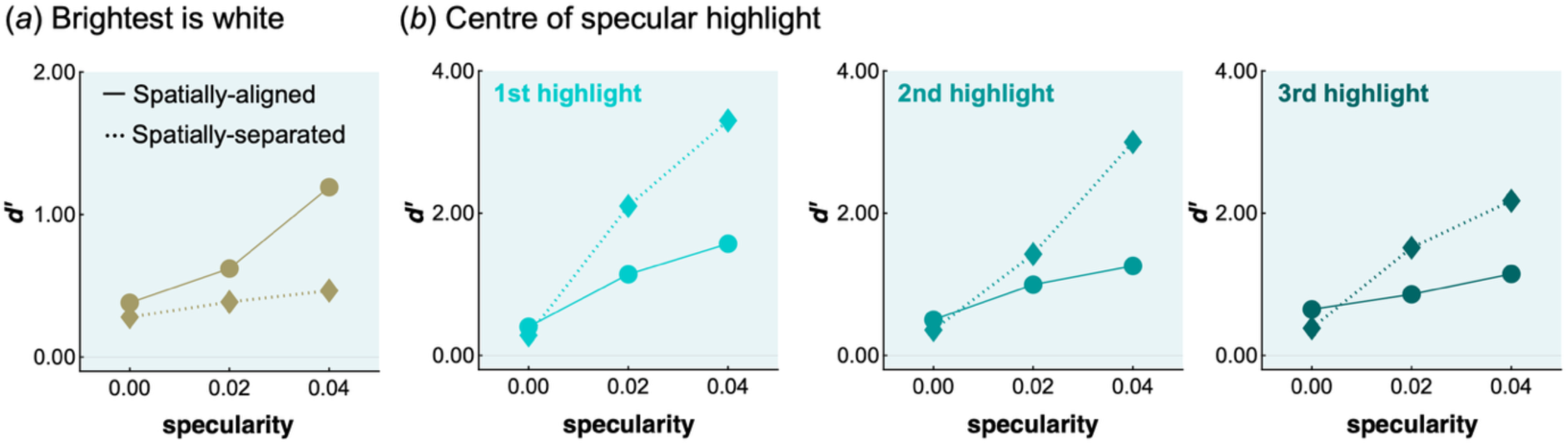
Patterns of discrimination sensitivity (*d’*) for two competing models. (*a*) The ‘brightest element’ heuristic. (*b*) The centre of specular highlight’ model. The 1st, 2nd, and 3rd highlights represent the most intense, second most intense, and third most intense highlights, respectively. Note that the y-axis scale differs between panels (*a*) and (*b*).

To quantify this difference, we computed for each observer the Pearson correlation between human and model d′ across the six conditions, and compared the four computational observers across the six human observers using a Friedman repeated-measures test with follow-up paired t-tests on Fisher z-transformed correlations. To correct for the six pairwise comparisons, significance was assessed against a Bonferroni-adjusted threshold of *α* = 0.05/6 = 0.00833. The brightest-element model correlated only weakly with human performance (mean *r* = 0.34, s.d. = 0.11), whereas all three centre-of-highlight models correlated strongly (1st: *r* = 0.95, s.d.= 0.02; 2nd: *r* = 0.92, s.d.= 0.06; 3rd: *r* = 0.93, s.d.= 0.02). This difference was reliable across observers (Friedman test, *p* = 0.004): each centre-of-highlight model fitted human d′ significantly better than the brightest-element model (brightest vs 1st: *t*(5) = −14.95, *p* < 0.001; vs 2nd: *t*(5) = −5.31, *p* = 0.003; vs 3rd: *t*(5) = −14.7, *p* < 0.001). The three centre-of-highlight models were broadly equivalent: the 1st and 3rd models differed (*t*(5) = 7.25, *p* < 0.001), whereas the 1st vs 2nd (*t*(5) = 0.03, *p* = 0.975) and 2nd vs 3rd (*t*(5) = 0.79, *p* = 0.466) comparisons did not. It should be noted that each correlation was computed across six conditions, two of which (the matte conditions) lie at chance for both humans and models; these shared chance points impose a baseline correlation. It is notable that the brightest-element and centre-of-highlight models nonetheless diverge even under this constraint.

## 4. Discussion

### (a) Summary of findings

The present study showed that, when an alternative decision rule is more informative, human observers can override the “brightest is white” heuristic. This suggests that the dependence of colour constancy on this heuristic is more limited, and more context-dependent, than its prevalence might imply, consistent with previous studies [22, 56]. When the brightest pixel was not the most reliable cue to illuminant colour, human observers’ performance improved, whereas a simulated ‘brightest element’ heuristic predicted a decline. This improvement can be attributed to specular highlights appearing on darker textured areas, where they provide a more accurate estimate of illumination colour due to reduced contamination from the diffuse component. These findings suggest that human observers do not rely solely on the brightest point to estimate an illuminant colour. Instead, an ideal observer model based on the centre of specular highlights successfully predicted the performance differences between spatially-aligned and spatially-separated conditions. Additionally, in the phase-scrambled condition, human *d’* values dropped significantly, highlighting the importance of the geometric relationship between the illuminant and surface. Taken together, our results suggest that human observers rely on properties of specular reflections to infer illumination colour, overriding the more-simplistic and traditionally-assumed heuristic that estimates illuminant colour from the brightest element in a scene.

### (b) Limits of pure colour-statistics strategies and the role of geometry

It was shown that human performance dropped significantly under the phase-scrambled condition where overall statistical properties were preserved. This suggests that observers do not rely solely on raw chromatic/luminance statistics but instead identify regions of interest based on the geometric configuration of surfaces and illumination and the regularities of diffuse and specular components before extracting statistical information. These findings highlight the nature of colour constancy and object colour judgments, emphasising the role of geometric information and spatio-chromatic properties in material perception. This has important implications for computational vision, as many existing computer vision systems rely on statistical heuristics for illuminant estimation. Developing models that incorporate scene context and semantic meaning could substantially improve the accuracy of these algorithms [57, 58]

### (c) Why the specular-centre model outperformed humans

The performance based on the centre of the specular highlight exceeded that observed in human participants. Several factors may help account for this discrepancy. First, with ten distinct lighting positions randomly assigned to each stimulus, observers were required to detect specular highlights and monitor colour changes within a brief 1.5-second animation. Second, sensory noise in the colour estimation, or a temporal-integration process that incorporates intermediate, less informative frames, may have further reduced human performance. While incorporating these factors into the model would likely lower its *d’* values to better align with human observers, our primary goal was not to match absolute performance levels but to predict patterns of performance across conditions based on candidate cues.

### (d) Limitations of the study and future extension

Our stimuli do not capture all the cues available in real-world settings. The paradigm deliberately isolates specular-highlight geometry by removing many cues that support colour constancy in natural scenes—a rich surround, mutual illumination, multiple objects, cast shadows and explicit illuminant cues. This is a strength, as it allows the simple brightest-is-white heuristic to be tested against a highlight-geometry strategy. However, assessing the degree of generalization to scenes with multiple objects of varied shapes and materials is an important next step. A related question is whether more spatially complex, naturalistic lighting environments improve the visual system’s ability to maintain colour and lightness constancy, such that observers are less reliant on a particular source of information [59–63]. Such research represents a step toward understanding the full capabilities of human colour perception in the context of complex visual environments. From an applied perspective, the finding suggests that computer-vision white-balance and colour-constancy algorithms could be improved by attending to where highlights fall, not merely how brightly they shine.

### (e) Everyday colour constancy and the need for realistic stimuli

When clouds block sunlight, altering the brightness and colour of a scene, or when checking how well ingredients are cooked based on their appearance, we skilfully distinguish whether the change in reflected light is due to a shift in illumination colour or a change in the object’s colour itself. A substantial body of research has explored mechanisms underlying colour constancy [12, 13]. However, many studies have relied on simple stimuli. Fully elucidating the mechanisms of human colour constancy will require incorporating material properties that the visual system routinely encounters, such as specular reflections and surface textures. Building on a growing body of work that has moved towards more naturalistic stimuli, the specific contribution of the present study is to show that, when the brightest element is made unreliable in an operational colour constancy task, observers exploit the geometry of specular highlights rather than relying on the brightest-is-white heuristic.

### (f) Conclusion

In conclusion, we have shown that when the brightest element is an unreliable cue to the illuminant, human observers instead exploit the geometry of specular highlights to estimate illuminant colour. This suggests that specular highlights serve not only as cues to surface gloss but also as informative samples of the light source, and that illuminant estimation can draw on the surface geometry rather than on image intensity alone. More broadly, our findings reinforce the idea that colour constancy is supported by multiple complementary cues, with observers able to rely on whichever remains informative in a given scene.

## Acknowledgements

This project has received funding under UKRI-Wellcome Physics of Life (EP/W023873/1; PhysFEM). TM was supported by a Sir Henry Wellcome Postdoctoral Fellowship from Wellcome Trust (218657/Z/19/Z). For the purpose of open access, the author has applied a CC BY public copyright license to any Author Accepted Manuscript version arising from this submission.

## Author contribution

Conceptualization: TM, HES, RJL

Data curation: TM

Formal Analysis: TM

Funding acquisition: TM, HES

Investigation: TM

Methodology: TM, HES, RJL

Project administration: TM, HES

Resources: TM, HES, RJL

Software: TM, RJL

Supervision: HES

Validation: TM

Visualization: TM

Writing – original draft - TM

Writing – review & editing - TM, RJL, HES

## Supplementary Information

### Rationale for the limitations of the “brightest is white” assumption

We provide a detailed explanation on why information extracted using the “brightest is white” heuristic can be less informative than that obtained from the centre of a specular highlight. A glossy object exhibits both matte and specular reflections, and the specular component alone offers a precise representation of the illuminant’s chromaticity. However, equation (1) based on the Ward reflectance model [1] shows that at any point in the image, the degree of dominance of the specular component is determined by the position-dependent coefficients α(*x*,*y*) and β(*x*,*y*).

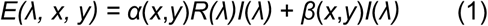

Here, *α*(*x*,*y*) and *β*(*x*,*y*) indicate coefficients due to scalar variation in the object’s surface reflectance (achromatic texture) and the magnitude of the specular component, respectively. R(*λ*) is the surface spectral reflectance of the sphere, and I(*λ*) is the illuminant spectral composition (sunlight or skylight). Thus, the first term in equation (1) corresponds to the diffuse reflection and the second term is the specular reflection. The scalar values α(x,y) varied between 0.2 (dark) and 1.0 (bright), and thus even in the dark part of the texture, the specular component and the diffuse component were always mixed, meaning that no direct information about the illuminant colour was presented in any region. Note that in addition to these factors, the intensity of the diffuse component is modified by a geometrical factor of the scene (i.e. the viewpoint, and the shape of the object) as the cosine of the angle between the surface normal and the light source. The specularity (a rendering parameter assigned to the surface) was constant over the whole surface, but the strength of specular reflection reaching the camera (i.e. in the image of the object) spatially varied due to the 3D geometry of the light source, surfaces and camera, and is strongest when the surface normal points mid-way between the source direction and camera.

In Figure 1*b* in the main text (spatially-separated condition), the “locus of chromaticities” over the whole object is represented by a black solid line. Because, at any point, the chromaticity is determined simply by the balance between diffuse and specular components, the whole chromatic locus is defined by the straight line between the plus and cross symbols. The reflected light from the location indicated by the green square has the furthest chromaticity from the illuminant because it does not contain any specular component. The reflected light from the cyan circle provides particularly good information about the illuminant because *α* is small and *β* is maximal at the centre of the highlight. The point indicated with a magenta diamond is less informative due to the higher *α* and smaller *β*. However, this is also the point selected by the “brightest element” heuristic, since the overall intensity at this location is higher than the intensity at the centre of highlight shown by the blue circle. Thus, for spatially-separated conditions, an optimal strategy is to use the centre of the specular highlight rather than a “brightest point” over the surface.

### The nature of the experimental task

In order to understand why each model resulted in the performances shown in Figure 4, it is helpful to understand the nature of the task and how the models can solve it. Figure 2*c* in the main text shows how the chromaticities of key points in the image change during the 1.5-seconds animation for an illuminant change (left) and a reflectance change (right) for specific trials. For an illuminant change, the chromatic distribution of the first frame and the final frame extends from their diffuse components to different illuminant chromaticities. Thus, if we focus on the centre of the specular highlight (cyan circle symbol) during the animation, the chromatic difference we observe should be close to the distance between the two illuminant chromaticities. On the other hand, a reflectance change shows a different change in chromatic distribution as each distribution is extending towards the same illuminant chromaticity. Thus, the chromatic difference in the centre of the highlight would become small. Therefore, the key to solving this task is to select a region that gives information about the illumination. It is worth emphasising that if we instead pick a point that has a diffuse component only, the chromatic change will be similar between the reflectance change and illuminant change trials due to the experimental design.

Figure S1 presents histograms of the chromatic difference across all trials for two model selection strategies: (*a*) choosing the brightest pixel and (*b*) selecting the centre of the most intense specular highlight. The leftmost panels depict learned distributions, based on 300 trials (50 reflectance changes and 50 illuminant changes for each specular level). For (a) the brightest pixel, the peaks of the reflectance change and illuminant change distributions are close together. In contrast, for (*b*) the centre of the specular highlight, the two distributions are more separated. The black line marks the intersection of these distributions, representing the criterion line. When the observed difference exceeds this threshold, the model classifies the trial as an illuminant change. Thus, the classification accuracy depends on which side of the line the observed chromatic difference falls. The right-hand histograms break down the left distributions by condition. When specularity is 0.00, both models produce identical, overlapping histograms, as expected. For the brightest pixel model, in the spatially-aligned condition, distributions separate effectively at specularity 0.04, allowing strong model performance. However, in the spatially-separated condition, separation is weaker. Conversely, for (*b*) the centre of the specular highlight, the trend is different from the brightest pixel model, aligning with the *d’* patterns in Figure 4 in the main text. Notably, at specularity 0.04, the model nearly perfectly separates the two distributions for spatially separated trials.

**Figure S1:**
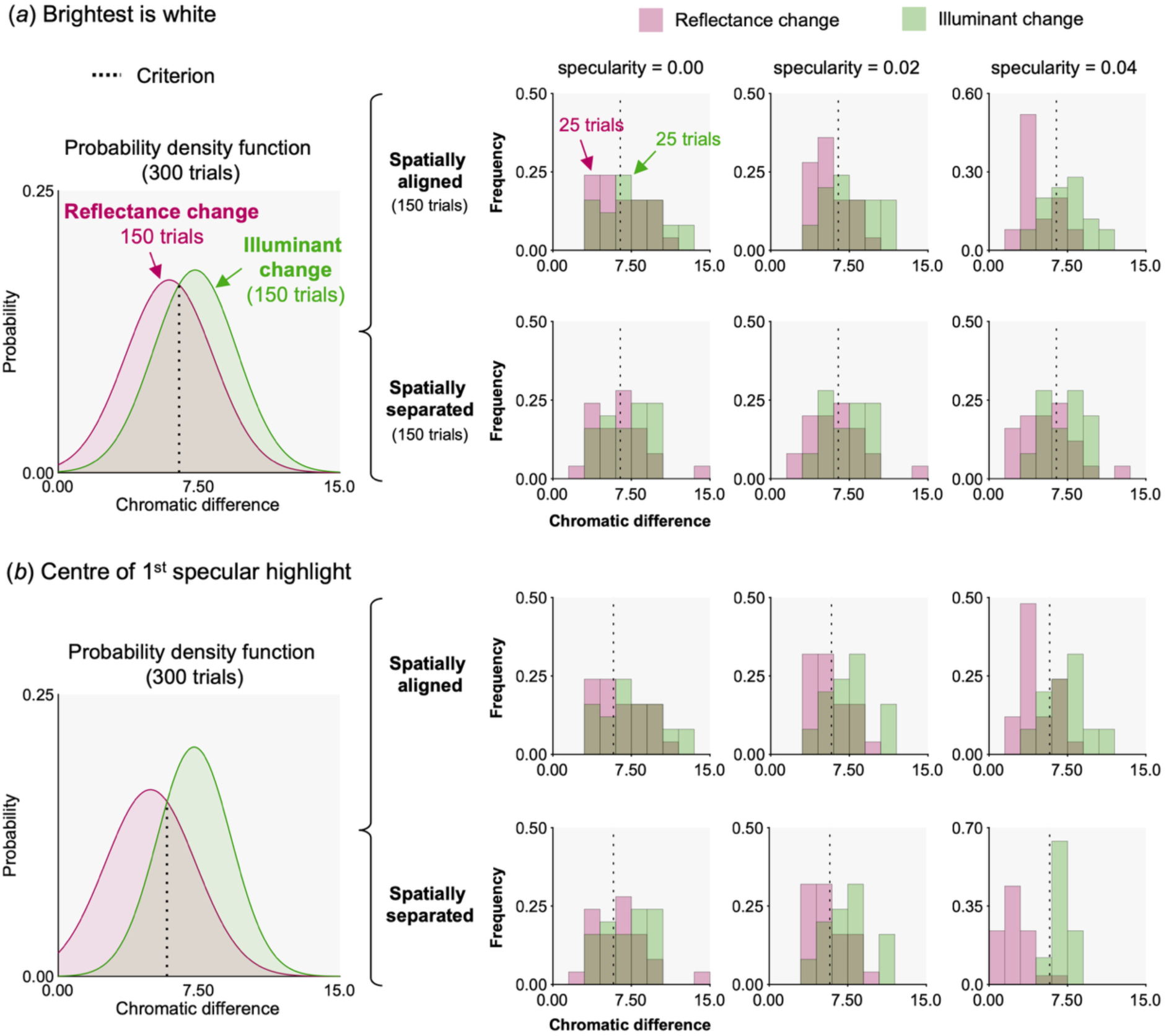
Histograms depicting the distribution of chromatic change magnitudes for (*a*) the brightest pixel and (*b*) the centre pixel of the specular highlight across all 300 trials (leftmost histogram) and for each condition (smaller histograms on the right). The curves in the leftmost histogram represent Gaussian fits to the data. The black dashed line shows the classification criterion, where chromatic differences smaller than this criterion are categorised as reflectance changes, and those larger are classified as illuminant changes.

### Using cone ratio to locate specular highlights

In the main text, the specular highlight model implicitly assumes that the location of the specular-highlight centre is known. But in practice, how can this position be estimated directly from the image signals themselves? To address this question, we simulated a simple detector that identifies specular-highlight regions by locating outliers in the cone-excitation ratios between the first and final frames. Our highlight-detection step was inspired by work on relational colour constancy, which showed that the spatial ratios of cone excitations between surface points remain approximately invariant under a change of illuminant [2, 3]. We adapted a similar idea to localise specular highlights. For matte points, reflected light is the product of reflectance and illuminant, so between the first and final frames the chromaticity of the diffuse surface should transform coherently, leaving its cone-excitation ratios largely preserved, whether the trial involves an illuminant or a reflectance change. At a specular highlight, a strong specular component is superimposed on this diffuse reflection. Because that component reflects the illuminant directly, it makes the highlight behave differently from the surrounding matte surface: adding an extra shift when the illuminant changes, and remaining comparatively fixed while the diffuse surround changes under a reflectance change. In either case, the specular contribution decouples the highlight’s cone-excitation ratios from those of the diffuse surface, so that pixels whose change in cone ratio between the first and final frames departs most from the global distribution are likely to coincide with specular highlights.

We analysed 100 stimuli (50 reflectance changes and 50 illuminant changes) at the highest specularity level. Each LMS image was blurred with a Gaussian filter (*σ* = 10 pixels) to suppress pixel-level noise. For the sphere stimuli, a circular crop was applied around the object to remove background regions and minimize edge artifacts. The same preprocessing steps were applied independently to frame 1 and frame 10.

For each cone class, we computed the pixel-wise ratio of responses between the two frames using equation (2):

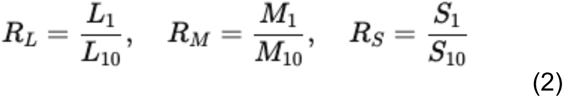

These ratios reflect the relative change in cone excitations between the two frames and form the basis for detecting local deviations indicative of specular highlights. To identify spatial outliers in cone-ratio statistics, each ratio map was standardized as in equation (3):

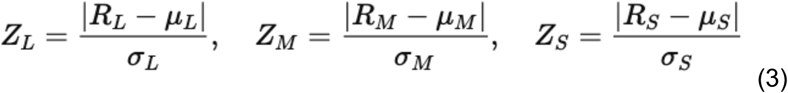

where *μ* and *σ* are the mean and standard deviation computed across all pixels.

A combined LMS deviation map was then computed using equation (4):

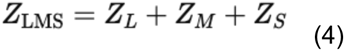

This provides a scalar measure of how strongly each pixel deviates from the global cone-ratio distribution; nine examples out of the 100 stimuli are shown in Figure S2. For the original sphere condition (panel *a*), for both spatially-aligned and spatially-separated configurations, specular reflections appear as clear outliers, allowing the specular-highlight centres to be identified straightforwardly once these regions are detected. In contrast, for the phase-scrambled condition (panel *b*), the specular pattern becomes spatially dispersed, and the highlight centre cannot be easily located. The detection was robust across the remaining 91 images, and repeating the analysis at the mid-specularity level yielded highly similar trends.

These results suggest that the statistical measure proposed in earlier work [2, 3] can also be applied effectively to the present stimuli, helping to separate the diffuse and specular components of the proximal image. We note, however, that this approach may be less effective for objects containing multiple diffuse reflectances, which can introduce substantial cone-signal variation across the surface.

**Figure S2:**
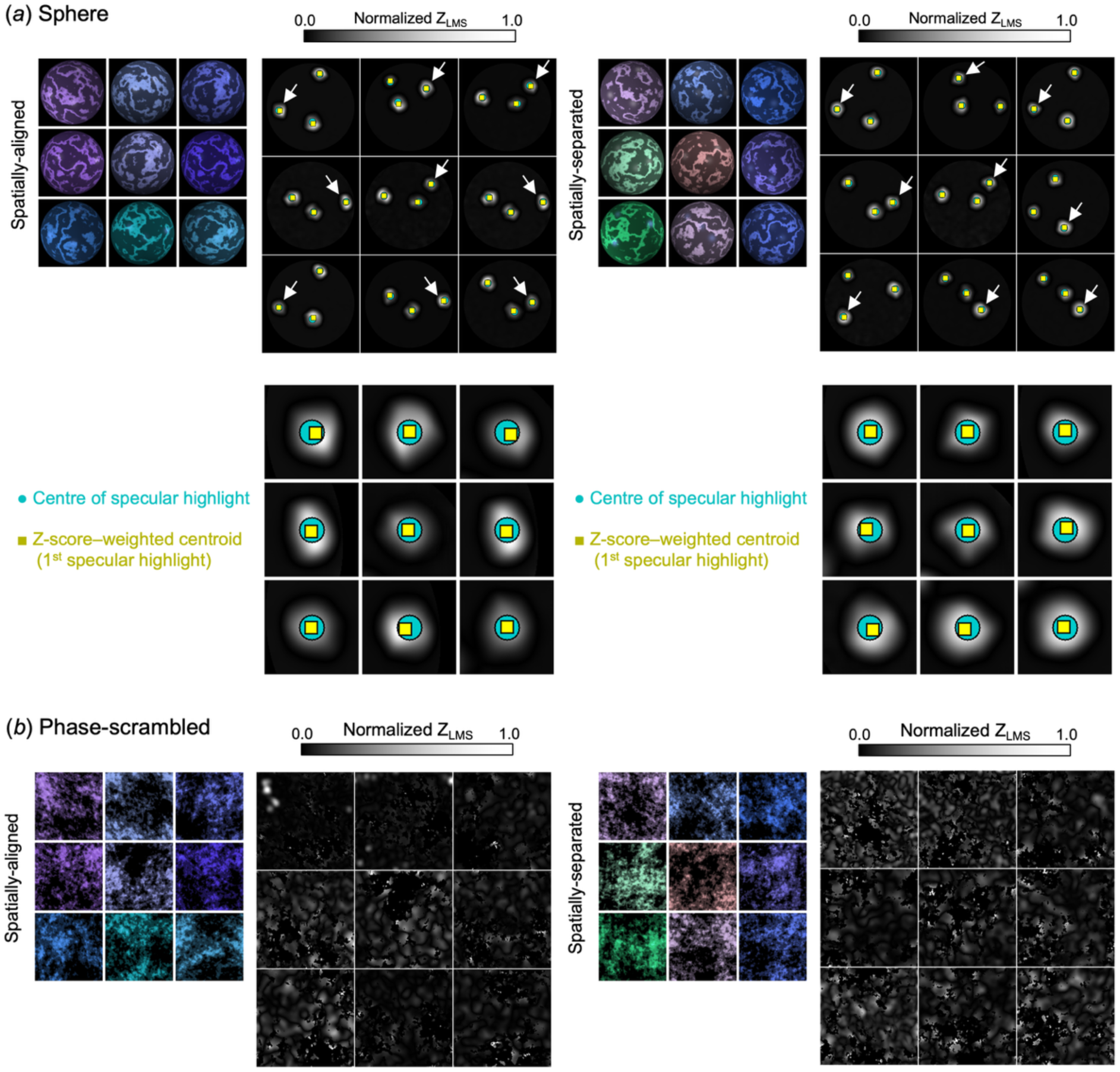
Detection of specular-highlight regions based on cone-ratio changes between the first and final frames. (*a*) Sphere stimuli. The left and right columns show the spatially aligned and spatially separated conditions, respectively. The top left panels display nine example RGB images (first frame only), and the top right panels show the corresponding Z_LMS_ maps, each normalized so that the 99.9th percentile equals 1. The bottom panels show cropped regions centred on the strongest specular highlight, indicated by the white arrow in the upper heatmaps. Colored symbols denote the centre of the specular highlight and the Z-score-weighted centroid around the first specular component, which closely aligns with the specular-highlight centre. The left and right columns correspond to the spatially-aligned and spatially-separated condition, respectively. (*b*) The same analysis applied to phase-scrambled versions of the images.

### Impact of spatial summation area around the point of interest

The spatial summation area for the visual system’s extraction of these statistics may extend beyond a single pixel in the image. To account for this, we implemented an alternative version of the computational model that averages chromaticities across different spatial scales around either the brightest pixel or the centre of the specular highlight—specifically, 3×3, 5×5, and 10×10 pixel regions. However, as shown in Figure S3, varying the spatial scale had minimal impact on the results.

**Figure S3:**
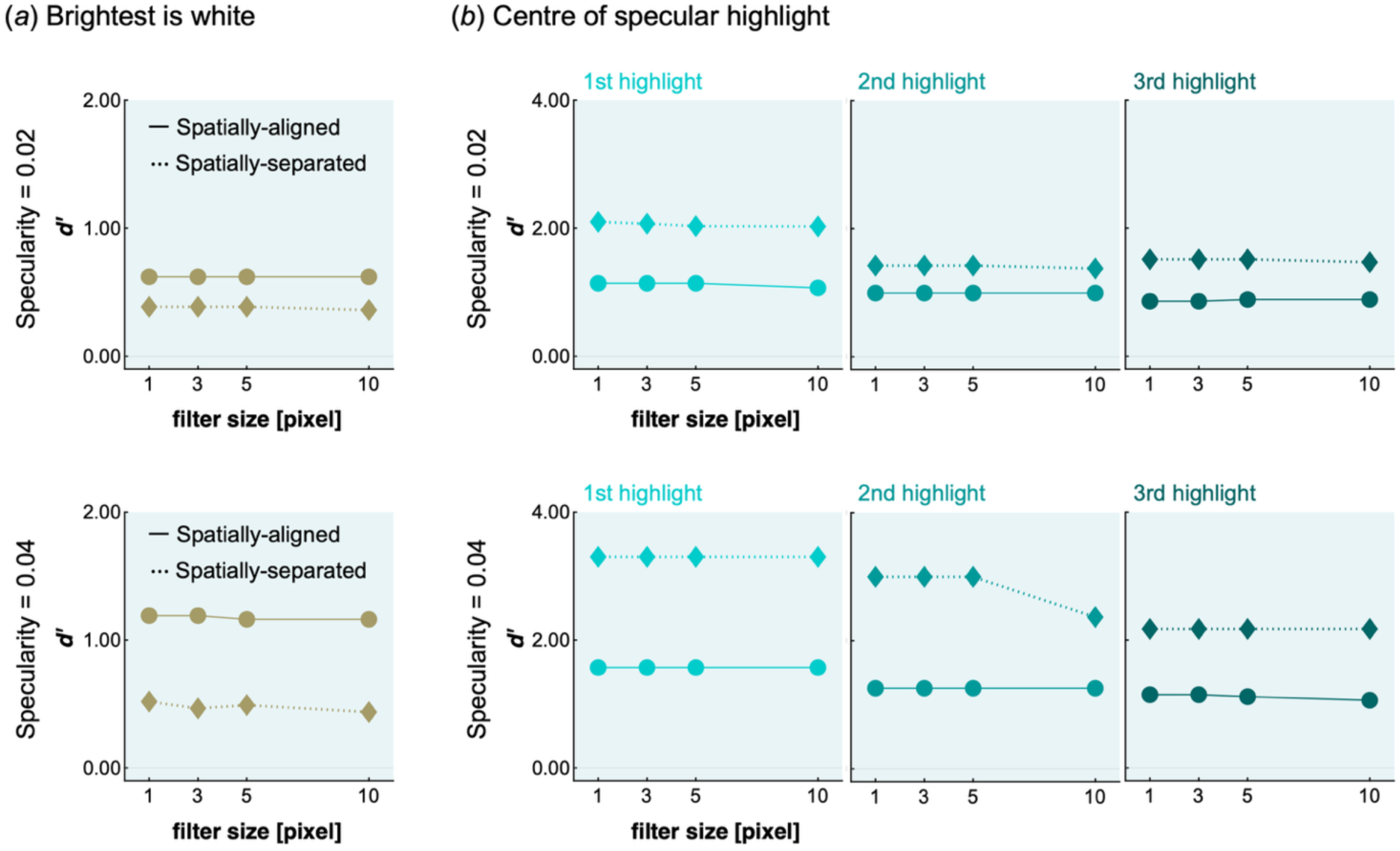
*d’* values computed when computational observers average over a square region of pixels of various sizes—one pixel (as reported in the main text), 3×3, 5×5, and 10×10 pixels—centred on the point of interest : (*a*) the brightest pixel and (*b*) the centre pixel of the specular highlight. The upper row corresponds to a specularity of 0.02, while the lower row corresponds to a specularity of 0.04.

